# Value, confidence, deliberation: a functional partition of the medial prefrontal cortex demonstrated across rating and choice tasks

**DOI:** 10.1101/2020.09.17.301291

**Authors:** N. Clairis, M. Pessiglione

## Abstract

Deciding about courses of action involves minimizing costs and maximizing benefits. Decision neuroscience studies have implicated both the ventral and dorsal medial prefrontal cortex (vmPFC and dmPFC) in signaling goal value and action cost, but the precise functional role of these regions is still a matter of debate. Here, we suggest a more general functional partition that applies not only to decisions but also to judgments about goal value (expected reward) and action cost (expected effort). In this conceptual framework, cognitive representations related to options (reward value and effort cost) are dissociated from metacognitive representations (confidence and deliberation) related to solving the task (providing a judgment or making a choice). Thus, we used an original approach with the goal of identifying consistencies across several preference tasks, from likeability ratings to binary decisions involving both attribute integration and option comparison. FMRI results confirmed the vmPFC as a generic valuation system, its activity increasing with reward value and decreasing with effort cost. In contrast, more dorsal regions were not concerned with the valuation of options but with metacognitive variables, confidence level being reflected in mPFC activity and deliberation time in dmPFC activity. Thus, there was a dissociation between the effort attached to choice options (represented in the vmPFC) and the effort invested in deliberation (represented in the dmPFC), the latter being expressed in pupil dilation. More generally, assessing commonalities across preference tasks might help reaching a unified view of the neural mechanisms underlying the cost/benefit tradeoffs that drive human behavior.

## Introduction

Standard decision theory assumes that selecting a course of action can be reduced to maximizing a net value function, where expected benefits are discounted by expected costs. Numerous studies in decision neuroscience have implicated key regions of the medial prefrontal cortex in computing the net values of options during choice. While there is a general agreement for a functional dissociation between ventral and dorsal parts of the medial PFC (vmPFC, sometimes called medial OFC, versus dmPFC, sometimes called dACC), the specific roles of these subregions are still a matter of debate.

Some accounts insist on the opponency between costs and benefits (Pessiglione et al., 2018; Rangel and Hare, 2010): the vmPFC would estimate the expected reward while the dmPFC would estimate the expected effort (Bartra et al., 2013; Clithero and Rangel, 2014; Kurniawan et al., 2013; Skvortsova et al., 2014). However, this view has been challenged by representations of effort cost found in vmPFC activity and reward value in dmPFC activity (Aridan et al., 2019; Arulpragasam et al., 2018; Fouragnan et al., 2015; Gläscher et al., 2009; Hogan et al., 2019; Klein-Flugge et al., 2016; Lopez-Gamundi et al., 2021; Pisauro et al., 2017; Seaman et al., 2018; Westbrook et al., 2019). Other accounts insist on the comparison between options that occurs during choice and suggest that the two regions estimate decision values in opposite fashion (Boorman et al., 2009; Hunt et al., 2012; Jocham et al., 2012; Wunderlich et al., 2009): the vmPFC would activate while the dmPFC would deactivate with value difference (chosen minus unchosen option value). Yet this other view has been questioned because the correlation with chosen and unchosen option values is not always observed in these regions and because the value difference may be confounded with other constructs such as default preference, choice confidence and decision time (Bobadilla-Suarez et al., 2020; De Martino et al., 2013; Jocham et al., 2014; Lim et al., 2011; Lopez-Persem et al., 2016; Massar et al., 2015; Qin et al., 2011). Thus, both types of accounts have received empirical support but also contradictory evidence, such that their validity is still debated.

Here, we intend to take a step aside from these debates and propose a functional partition that would generalize beyond choice tasks. Indeed, contrary to the view that there is no value representation outside of choice contexts (Hayden and Niv, 2021), neural correlates of values in the medial PFC have been found in many tasks that do not involve any choice between the items presented, including likeability rating and distractive tasks or passive viewing during which covert likeability ratings are spontaneously generated (Lebreton et al., 2009; Plassmann et al., 2010; Harvey et al., 2010; Levy et al., 2011; Abitbol et al., 2015; De Martino et al., 2017; Shenhav and Karmarkar, 2019; Lopez-Persem et al., 2020). We therefore reasoned that a general account for the role of the medial PFC in expressing preference should explain the pattern of activity observed during both rating and choice.

The new functional partition that we propose here is based on a metacognitive account (Lee and Daunizeau, 2021): the idea is that, whatever the task, the brain invests effort in deliberation until it reaches a satisfactory level of confidence in the intended response. Thus, a second cost/benefit tradeoff would govern the meta-decision about when to make a response, the cost being the amount of time spent in deliberation and the benefit being the level of confidence attained. During this double cost/benefit arbitration, the brain would represent two sort of variables: 1) at the decisional level, the reward and effort values associated to options proposed for rating or choice, and 2) at the metacognitive level, the expected confidence in the response and the required amount of deliberation. The aim of the present study is to test whether this functional partition can account for the pattern of activity observed in medial prefrontal regions across rating and choice tasks.

To this aim, we reversed the typical logic of standard functional neuroimaging approach, which specifies the roles of brain regions with contrasts that isolate minimal differences between conditions. On the contrary, we intended to generalize our findings across various conditions and tasks, with the aim to reach more robust conclusions. Thus, we employed a series of preference tasks (also called ‘value-based’ tasks) that enable the investigation of 1) the assignment of reward value or effort cost to a single option, with likeability rating tasks, 2) the comparison between two reward or two effort options with A/B choice tasks, and 3) the integration of reward and effort attributes for one option to accept or reject, with Yes/No choice tasks. In all these tasks, we defined the same key variables of interest as the global stimulus value (Val), which increases with more appetitive reward and/or less aversive effort, the confidence in the response (Conf), which is higher for more extreme ratings and more likely choices, and deliberation time (DT), meaning duration of the effort invested in the valuation process so as to reach a satisfactory response. We then explored the relationships between these three variables at the behavioral level, and their representations in the medial PFC at the neural level.

## Results

### Behavior

Participants (n=39 in total, 22 females) first performed a series of ratings, divided into three fMRI sessions (Fig. 1A). Each session presented 72 items to be valuated one by one. Within a session, items were grouped into three blocks: one block with 24 reward items presented by text + image (R_ti_), one block with 24 reward items presented by text only (R_t_) and one block with 24 effort items presented by text only (E_t_). The reason for varying the mode of presentation was to assess the generality of the neural valuation process across different inputs that require more or less imagination, according to previous study (Lebreton et al., 2013). For reward, participants were asked to rate how much they would like it, should they be given the item immediately after the experiment. Symmetrically, the instruction for effort was to rate how much they would dislike it, should they be requested to exert it immediately after the experiment. We included both food and non-food (goodies) reward items, and both mental and physical effort items. There was no number on the scale, just labels on endpoints, and ratings were pseudo-continuous, from ‘I would not care / mind’ to ‘I would like / dislike it enormously’. Thus, the left endpoint corresponded to indifference and the right endpoint to extreme attraction or extreme aversion (Fig. 1A).

**Figure 1.**
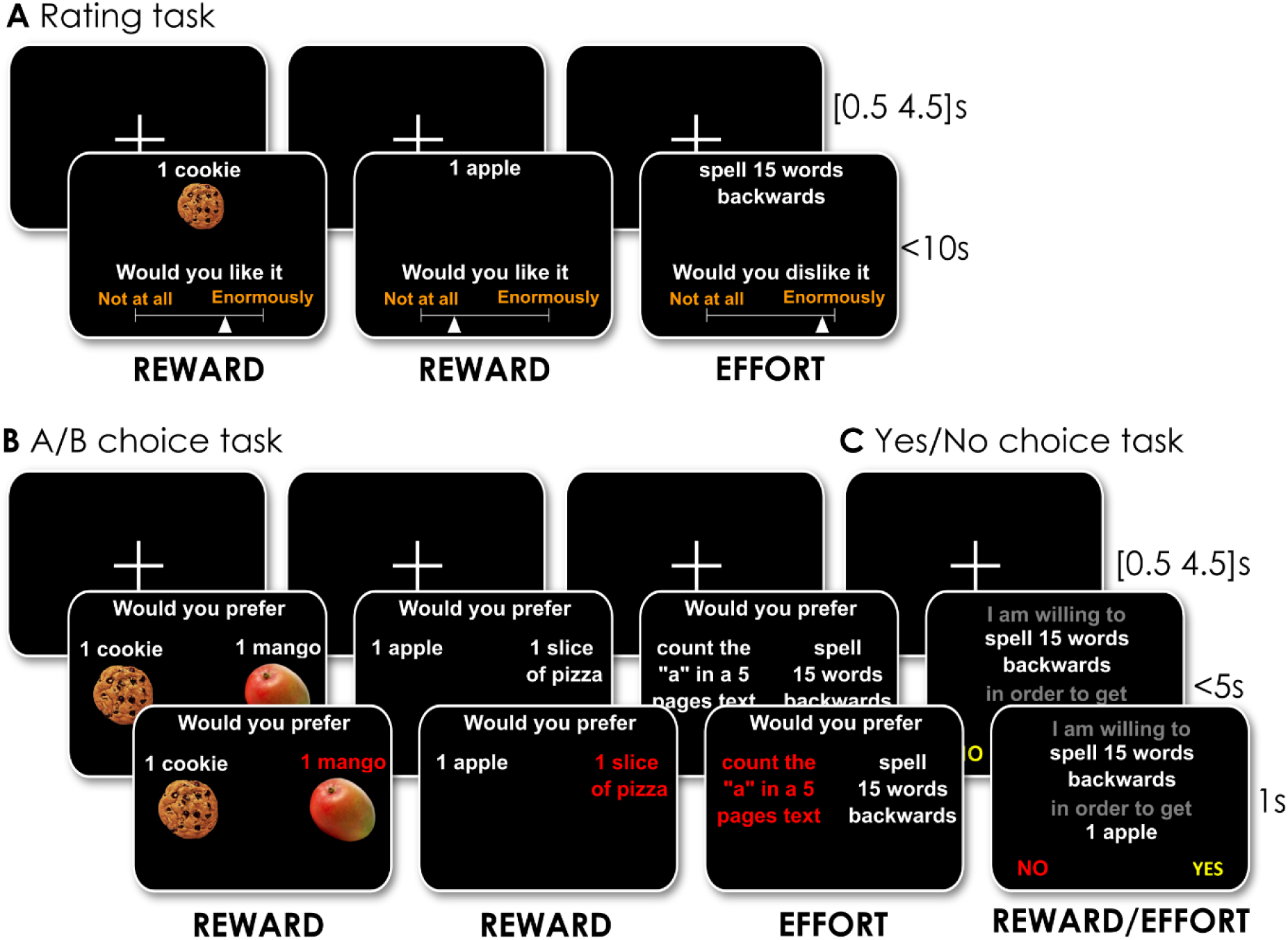
Behavioral tasks. Example trials are illustrated as a succession of screenshots from top to bottom, with durations in seconds. Only the duration of fixation cross display at the beginning of trials is jittered. The duration of the response screen depends on deliberation time, as both rating and choice are self-paced. A] Rating task. In every trial, subjects are shown an item that can be a reward described with both text and image (R_ti_), a reward described with text only (R_t_) or an effort described with text only (E_t_). The task for subjects is to rate how much they would like receiving the proposed reward or dislike performing the proposed effort, should it occur, hypothetically, at the end of the experiment. They first move the cursor using left and right buttons on a pad to the position that best reflect their (dis)likeability estimate, then validate their response with a third button and proceed to the next trial. B] A/B choice task. In every trial, two options belonging to the same category are shown on screen and subjects are asked to select their favorite option, i.e. which reward they would prefer to receive if they were offered the two options or which effort they would prefer to exert if they were forced to implement one of the two options at the end of the experiment (hypothetically). The choice is expressed by selecting between left and right buttons with the index or middle finger. The chosen option is then highlighted in red, and subjects proceed to the next trial. C] Yes/No choice task. In every trial, one option combining the two dimensions is shown on screen and subjects are asked to state whether they would be willing to exert the effort in order to receive the reward, if they were given the opportunity at the end of the experiment (hypothetically). They select their response (‘yes’ or ‘no’, positions counterbalanced across trials) by pressing the left or right button, with their index or middle finger.

The z-scored rating was taken as a proxy for stimulus value (Val) in this task, while the square of z-score rating was taken as a proxy for response confidence (Conf). The quadratic relationship between confidence and rating has been validated empirically and accounted for by a Bayes-optimal model mapping a probabilistic distribution (over likeability) onto a bounded visual scale (Lebreton et al., 2015; Lopez-Persem et al., 2020). Under this model, confidence is inversely proportional to the variance of the underlying probability distribution, hence to the variability in likeability rating across presentations of the same item when they are repeated (which was not the case in the present design). The confidence proxy used here is not to be confounded with motivational salience, which would be maximal for very appetitive reward and very aversive effort. Instead, confidence is maximal at the extremes of the rating scale, meaning for both very appetitive and null reward or for both very aversive and null effort (Fig. 2A). Note also that Val and Conf were orthogonal variables by construction (Conf being a U-shaped function of Val for both reward and effort).

**Figure 2:**
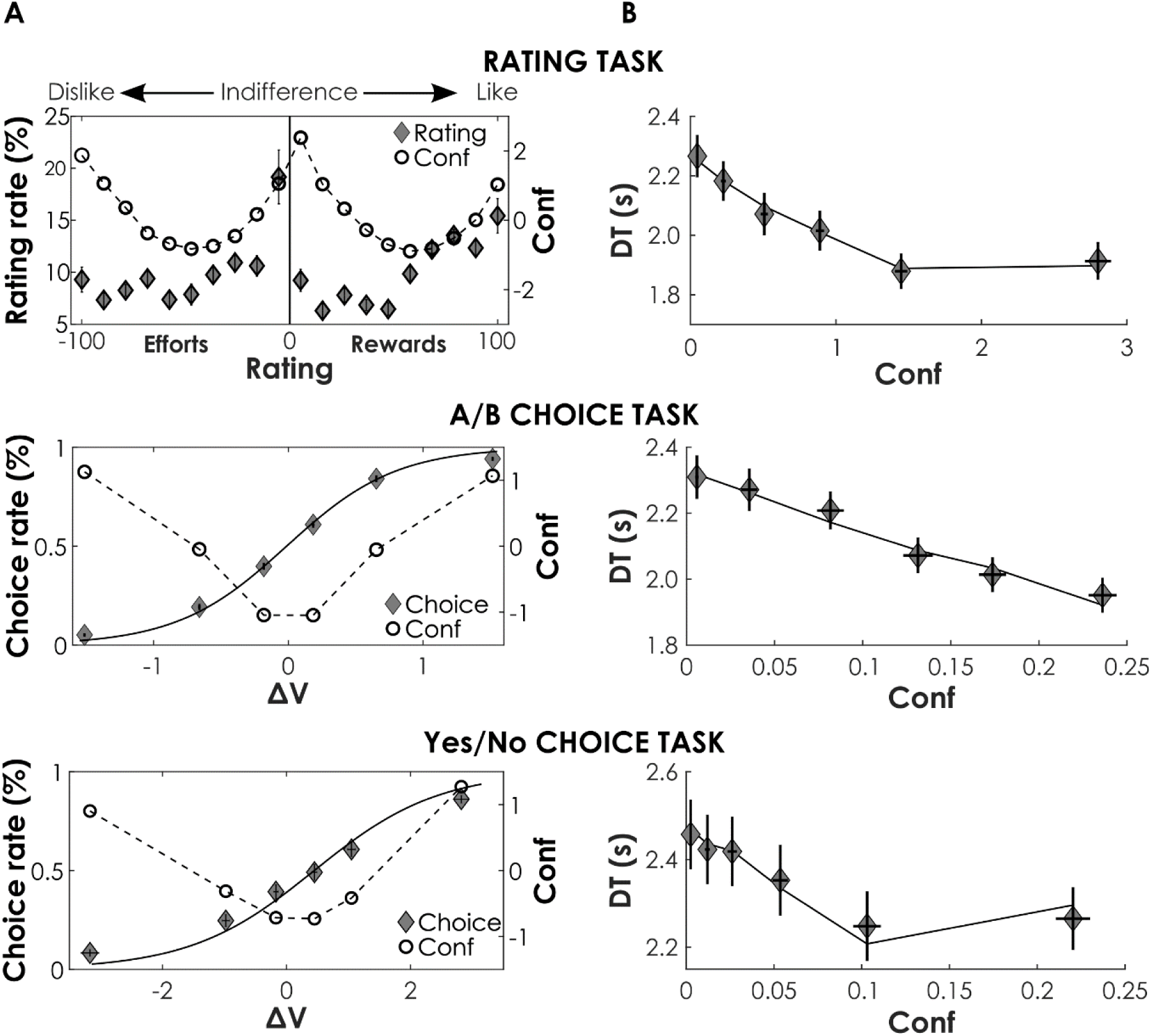
Behavioral results. A] Response rate. For ratings, plots show the average response rate for each bin (portion of the rating scale). Effort items (on the left) are rated between bin 0 (‘I would not mind’) and bin -10 (‘I would dislike it enormously’). Reward items (on the right) are rated between bin 0 (‘I would not care’) and bin +10 (‘I would like it enormously’). Note that the x-axis has been reverted for effort ratings, compared to the visual scale presented in the task, such that it globally indicates increasing values (less aversive effort from -100 to 0 and more appetitive rewards from 0 to +100). For choices, the response rate is plotted as a function of binned decision value (ΔV). In the A /B task, decision value is the difference in likeability rating between left and right options (V_left_ − V_right_), and choice rate is the frequency of left option being selected. In the Yes/No task, decision value is the addition of weighted reward and effort likeability ratings (β_R_·V_R_ + β_E_·V_E_), which is equivalent to both stimulus value (Val) and to the value difference between yes and no options (net value minus zero). Continuous lines show logistic regression fits of choice rate and dashed lines show variations in the confidence proxy (Conf). B] Deliberation time as a function of confidence proxy (Conf), defined as the square of centered likeability rating (V^2^) for rating tasks and the square of centered choice likelihood (P^2^) for choice tasks. The Conf proxy was validated in two different datasets where confidence in rating or choice was directly asked to participants (see Fig. S1A). Dots represent mean across participants, x and y error bars are inter-participant standard errors.

Deliberation time (DT) was defined as the time between item onset and the first button press used to move the cursor along the scale. DT was regressed against a linear model that included Val and Conf proxies (Fig. 2B), in addition to factors of no interest (such as jitter duration, stimulus luminance, text length and trial index, see methods). Irrespective of stimulus type, we found a significant effect of both value (R_ti_: β_Val_ = -0.21 ± 0.02, p = 4·10^−11^; R_t_: β_Val_ = -0.17 ± 0.02, p = 6·10^−11^; E_t_: β_Val_ = 0.26 ± 0.03, p = 2·10^−11^) and confidence (R_ti_: β_Conf_ = -0.17 ± 0.03, p = 3·10^−8^; R_t_: β_Conf_ = -0.19 ± 0.03, p = 7·10^−8^; E_t_: β_Conf_ = -0.13 ± 0.04; p = 0.0024). Thus, participants were faster to provide their rating when the item was more appetitive (or less aversive) and when they were more confident (going towards the extremes of the rating scale). Among the factors of no interest, we observed effects of jitter duration, stimulus luminance and text length, which were therefore included as regressors in subsequent analyses. However, there was no significant effect of trial index, which discards a possible contamination of DT by habituation or fatigue.

Then participants performed a series of binary choices, either A/B choices or Yes/No choices. The choice tasks were always performed after the rating tasks because the ratings were used to control the difficulty of choices (i.e., the difference in value between the two options). In the A/B choice task (Fig. 1B), participants were asked to select the reward they would prefer to receive at the end of the experiment, if they were offered one of two options, or the effort they would prefer to exert, if they were forced to implement one of two options. Thus, the two options always pertained to the same dimension (reward or effort), and even to the same sub-category (food or good for reward, mental or physical for effort), to avoid shortcut of deliberation by general preference. The mode of presentation (text or image) was also the same for the two options, to avoid biasing the choice by a difference in salience. To obtain a same number of trials as in the rating task, each item was presented twice, for a total of 72 choices per stimulus type (R_ti_, R_t_, E_t_) distributed over three fMRI sessions. Within a session, items were grouped into three blocks: one block with 24 choices between reward items presented with text + image (R_ti_), one block with 24 choices between reward items presented with text only (R_t_) and one block with 24 choices between effort items presented with text only (E_t_). In the Yes/No choice task (Fig. 1C), participants were asked whether they would be willing to exert an effort in order to obtain a reward, at the end of the experiment. Only items described with text were retained for this task (since there was no picture for effort items), each item again appearing twice, for a total of 144 choices divided into three fMRI sessions of 48 trials each.

The A/B choice task was meant to assess value comparison between the two options, within a same dimension. The decision value (ΔV) in this task was defined as the difference in (dis-)likeability rating between the two options. We checked with a logistic regression (Fig. 2A) that ΔV was a significant predictor of choices, irrespective of stimulus type (R_ti_: β_ΔV_ = 3.38 ± 0.27, p = 7·10^−15^; R_t_: β_ΔV_ = 2.67 ± 0.16, p = 2·10^−19^; E_t_: β_ΔV_ = -2.28 ± 0.16, p = 4·10^−17^). The Yes/No choice task was meant to assess value integration across two dimensions, for a single option. The decision value (or net value) in this task was defined as a linear combination of reward and effort ratings. Note that it would make no sense to fit an effort discounting function here, because such function is meant to capture the mapping from objective effort levels to subjective effort estimates, which we directly collected (with dislikeability ratings). We checked with a logistic regression that both reward and effort ratings were significant predictors of choice in this task (β_R_ = 1.50 ± 0.09, p = 6·10^−20^; β_E_ = -1.12 ± 0.08, p = 1·10^−16^).

To analyze DT (time between stimulus onset and button press) in choice tasks, we defined proxies for stimulus value and response confidence, as we did for the rating task. Stimulus value (Val) was defined as the addition of the likeability ratings assigned to the two stimuli on screen. In the A/B choice task, this is simply the sum of the two item ratings. In the Yes/No choice task, this is a weighted sum (with a scaling factor to adjust the unit of reward and effort ratings). In both cases, choice probability was calculated with the logistic regression model (softmax function of decision value). Response confidence (Conf) was defined, by analogy to the rating task, as the square of the difference between choice probability and mean choice rate. Linear regression showed that DT decreased with value in the A/B choice task (R_ti_: β_Val_ = -0.06 ± 0.01, p = 3·10^−7^; R_t_: β_Val_ = -0.06 ± 0.01, p = 3·10^−7^; E_t_: β_Val_ = 0.05 ± 0.01, p = 8·10^−4^), albeit not in the Yes/No choice task (β_Val_ = 0.033 ± 0.024, p = 0.172). DT also decreased with confidence (Fig. 2B) in both the A/B choice task (R_ti_: β_Conf_ = -1.74 ± 0.20, p = 2·10^−10^; R_t_: β_Conf_ = -1.98 ± 0.18, p = 4·10^−13^; E_t_: β_Conf_ = -1.73 ± 0.22, p = 2·10^−9^) and the Yes/No choice task (β_Conf_ = -1.15 ± 0.15, p = 1·10^−9^). Thus, the relationship between DT and confidence was similar in rating and choice tasks: participants were faster when they were more confident (because of a strong preference for one response or the other). They also tended to be faster when the options were more appetitive (or less aversive), but this trend was not significant in all tasks.

Because we did not measure confidence in the present study, we verified that our proxy could predict confidence ratings in separate datasets. Note that this proxy has already been validated for likeability rating tasks used in previous studies (De Martino et al., 2017; Lebreton et al., 2015; Lopez-Persem et al., 2020), a result that we reproduced here (Fig. S1B). To test whether the same proxy could also predict confidence in choice tasks, we used another dataset from a published study (Lee and Daunizeau, 2020). In this study (Fig. S1A), participants provided confidence ratings about having selected the best option in binary A/B choices (between food items presented two by two). Our confidence proxy could significantly predict confidence judgments not only in the likeability rating task but also in the A/B choice task even when including Val and DT as competitors (without orthogonalization) in the same regression model (rating: β_Conf_ = 0.49 ± 0.09; p = 8·10^−5^; choice: β_Conf_ = 0.21 ± 0.02; p = 2·10^−11^).

### Neural activity

The aim of fMRI data analysis was to dissociate the first-level variables related to option attributes (reward and effort estimates) from the second-level variables related to metacognition (confidence and deliberation) across value-based tasks (rating and choice). To assess whether these variables can be dissociated on the basis of existing literature, we conducted a meta-analysis of fMRI studies using Neurosynth platform (Fig. 3A) with value, confidence and effort as keywords. Results show that the three keywords are associated to similar activation patterns, with clusters in both vmPFC and dmPFC. To better dissociate the neural correlates of these constructs in our dataset, we built a general linear model where stimulus onset events were modulated by our three variables of interest - Val, Conf and DT (defined as in the behavioral data analysis). Factors of no interest that were found to influence DT (jitter duration, stimulus luminance, text length) were also included as modulators of stimulus onset events, before the variables of interest. Note that by construction, the correlation between regressors of interest was low (between -0.084 and -0.204). Nevertheless, to avoid any confound in the interpretation, we employed serial orthogonalization. Thus, the variables of interest were orthogonalized with respect to factors of no interest, and DT was made orthogonal to all other regressors, including Val and Conf.

**Figure 3:**
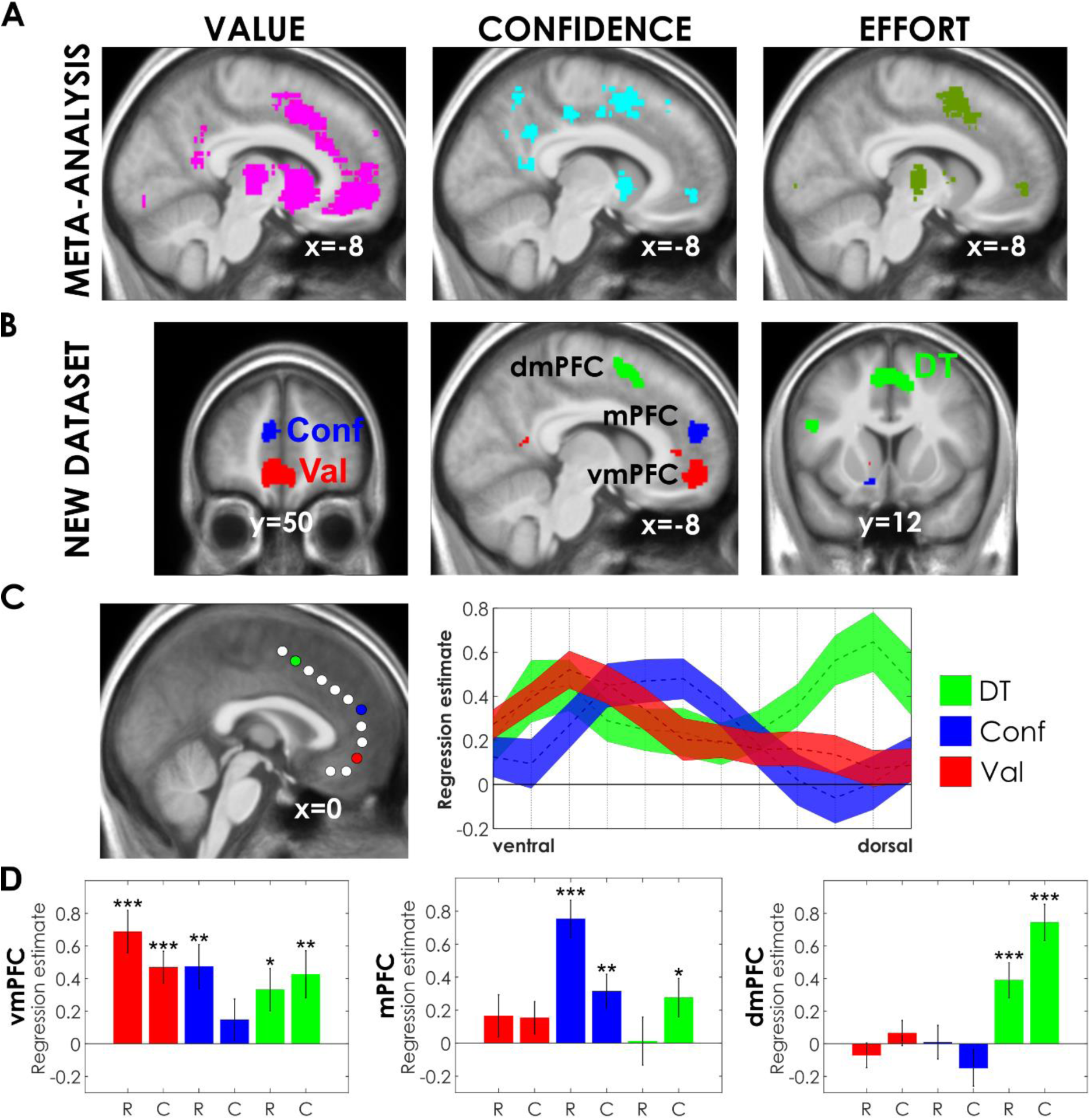
Neural results. A] Meta-analysis of fMRI studies. Statistical maps (sagittal slices) were extracted from the Neurosynth platform with the ‘value’, ‘confidence’ and ‘effort’ keywords. Significant clusters in the medial prefrontal cortex are similar across keywords, being located in both ventral and dorsal regions. B] Neural correlates of value, confidence and deliberation constructs in the present dataset. Statistical maps were obtained with a GLM including the different variables as parametric modulators of stimulus onset, across rating and choice tasks. Sagittal slice was taken at the same coordinates as the Neurosynth output, and superimposed on the average anatomical scan normalized to canonical (MNI) template. Coronal slices show the extent of the different medial prefrontal clusters. Statistical threshold was set at p < 0.05 after family-wise error for multiple comparisons at the voxel level. For clusters outside the medial prefrontal cortex, see activations in Tables S1, S2 and S3. For clusters obtained using the same GLM without orthogonalization of regressors and using the same GLM with events modeled as boxcar instead of stick functions, see Fig. S2 and Tables S4, S5 and S6. C] Distribution of regression estimates (inter-subject means ± standard errors) obtained for Val, Conf and DT variables along a ventro-dorsal line within the medial prefrontal cortex (sampled in each 8mm-radius shown on the average anatomical map). Colored circles show sampled spheres in which correlation with the corresponding variable was maximal (Val – red, Conf – blue and DT – green). D] Decomposition of regression estimates obtained for each variable of interest, plotted separately for rating and choice tasks (noted R and C) and for the different ROI (vmPFC, mPFC, dmPFC). Similar results were obtained when using different fMRI sequences (Fig. S3). For further investigations of value representation in the vmPFC, see Fig. S4. For the distinction between choice tasks (A/B *vs* Yes/ No), see Fig. S5. Bars show mean across participants; error bars show inter-participant standard errors. Stars indicate significance of t-test against zero (*** p < 0.001, ** p < 0.01, * p< 0.05).

After correction for multiple comparisons at the voxel level, we found only three significant clusters in the prefrontal cortex (Fig. 3B): Val was signaled in vmPFC activity (Table S1), Conf in mPFC activity (Table S2) and DT in dmPFC activity (Table S3). All three correlations were positive, there was no significantly negative correlation in any brain region when correcting for multiple comparisons. With a more lenient threshold (correction at the cluster level), we observed significant positive association with Val in other brain regions, such as the ventral striatum (vS), posterior cingulate cortex (pCC) and primary visual cortex (V1). Note that vS and pCC are standard components of the brain valuation system, whereas V1 activation is likely to be an artifact of gaze position on the rating scale, as it was not observed in the choice tasks. Consistently, positive correlation with Val was found in right V1 activity, and negative correlation in left V1 activity (a pattern that was not observed with other clusters). To provide a more exhaustive depiction, we examined the distribution of regression estimates below statistical thresholds, along a path going from vmPFC to dmPFC within a medial plane (Fig. 3C). Results show that the three associations did not correspond to separate clusters (as was suggested by thresholded maps) but to gradual variations peaking at different positions along the path.

To assess whether the triple association between variables and clusters of interest was robust, we conducted a number of additional analyses using variants of the main GLM (Fig. S2). The same three clusters were significantly associated with the Val, Conf and DT regressors when 1) removing serial orthogonalization such that regressors could compete for variance and 2) replacing stick functions by boxcar functions extending from stimulus onset to behavioral response (showing a modulation of dmPFC activity by DT in amplitude and not just duration). In addition, we tested the triple association using different fMRI acquisition sequences in participants of the pilot study (n=15). The fMRI sessions acquired with multiband acceleration sequences (see methods) were not included in the main analysis, since they were not directly comparable to those using our standard EPI sequence. We separately regressed fMRI activity recorded during these sessions against our main GLM, and observed similar trends in this independent dataset. Due to a three times smaller sample, activations did not pass whole-brain corrected thresholds. However, using group-level significant clusters (from the main dataset) as regions of interest (ROI), we observed significant associations of Val and DT with vmPFC and dmPFC, respectively (Fig. S3).

We further analyzed the relationship between computational variables and activity in the three medial prefrontal ROI with post-hoc t-tests on regression estimates. To avoid any double-dipping issue, we used a leave-one-out procedure, such that clusters were defined from group-level analyses including all subjects but the one in whom regression estimates were extracted. We first verified that the three main associations were not driven by any particular task (Fig. 3D). Indeed, regression estimates were significant in both rating and choice tasks, more specifically for Val in vmPFC activity (rating: β_Val_ = 0.69 ± 0.13, p = 6·10^−6^ ; choice: β_Val_= 0.47 ± 0.10, p = 3·10^−5^), for Conf in mPFC activity (rating: β_Conf_ = 0.75 ± 0.11, p = 8·10^−8^ ; choice: β_Conf_ = 0.31 ± 0.10, p = 0.004) and for DT in dmPFC activity (rating: β_DT_ = 0.39 ± 0.11, p = 9·10^−4^ ; choice: β_DT_ = 0.74 ± 0.11, p = 7·10^−8^). Note that our point was to generalize the associations across different tasks - comparing between tasks would be meaningless because tasks were not designed to be comparable (any possible significant contrast could be due to many differences of no interest).

We also investigated whether each cluster of interest was better associated with the corresponding variable (across tasks), again using a leave-one-out procedure to avoid double dipping (Fig. 3D): Val was better reflected in vmPFC activity (β_Val/vmPFC_ > β_Val/mPFC_ : p = 9·10^−8^ ; β_Val/vmPFC_ > β_Val/dmPFC_ : p = 4·10^−7^), Conf in mPFC activity (β_Conf/mPFC_ > β_Conf/vmPFC_ : p = 0.0043; β_Conf/mPFC_ > β_Conf/dmPFC_ : p = 3·10^−7^) and DT in dmPFC activity (β_DT/dmPFC_ > β_DT/vmPFC_ : p = 0.066; β_DT/dmPFC_ > β_DT/mPFC_ : p = 7·10^−4^). However, the fact that vmPFC, mPFC and dmPFC better reflected Val, Conf and DT, respectively, does not imply that these regions were not affected by the other variables. In particular, vmPFC activity was also associated with Conf and DT, (β_Conf_ = 0.26 ± 0.10, p = 0.012; β_DT_ = 0.40 ± 0.11, p = 0.001), even if it was dominated by Val-related activity. Nevertheless, all cross-over interactions between regions and variables were significant: from vmPFC to mPFC, the relative encoding of Val and Conf (β_Val_ - β_Conf_) significantly reversed (0.29 ± 0.11 vs. -0.30 ± 0.10, p=2·10^−8^) and similarly, from mPFC to dmPFC, the relative encoding of Conf and DT (β_Conf_ - β_DT_) significantly reversed (0.27 ± 0.13 vs. -0.72 ± 0.14, p = 9·10^−6^). The distant cross-over interaction between vmPFC and dmPFC (β_Val_ - β_DT_) was also significant (0.15 ± 0.15 vs. -0.30 ± 0.10, p=10^−5^).

We next looked for further generalization of the valuation signal, not solely across tasks but also across stimuli. In the main analysis, fMRI time series were regressed against a GLM that separated stimulus types (Rti, Rt and Et) into different onset regressors, each modulated by corresponding ratings. Instead of testing the average regression estimates across stimulus categories, we tested regression estimates obtained for each category, separately (Fig. S4A and S4B). Regression estimates (extracted using leave-one-out procedure across rating and choice tasks) show that vmPFC activity was positively related to the subjective value of reward items, whether or not they are presented with an image (Rti: β_Val_ = 0.49 ± 0.13, p = 8·10^−4^; Rt: β_Val_ = 0.61 ± 0.13, p = 5·10^−5^), and negatively correlated to the subjective cost of effort items (Et: β_Val_ = -0.35 ± 0.13, p = 0.017). Thus, the association between Val and vmPFC activity was independent of the presentation mode, and integrated costs as well as benefits.

On a different note, we questioned the validity of our Val proxy to capture value-related activity in choice tasks. Again, the reason for summing stimulus values in choice tasks instead of taking the difference between chosen and unchosen option values, as is often done, was that we wanted a proxy that could generalize to rating tasks, in which there is no notion of difference, since there is only one stimulus on screen. Note that the value difference regressor (chosen minus unchosen option value) is related to all three variables that we intend to dissociate here as capturing different concepts (stimulus value, response confidence, deliberation effort). Nevertheless, we wondered whether vmPFC activity in choice tasks would be better captured by the difference (V_c_ − V_uc_) than by the sum (V_c_ + V_uc_). To test this, we simply replaced our partition (Val / Conf / DT) by V_c_ and V_uc_ regressors, and fitted the GLM to fMRI activity recorded during choice tasks only (Fig. S4C). The two regression estimates, extracted from the Val cluster in the main analysis, were significantly positive (β_Vc_= 0.42 ± 0.12, p = 9·10^−4^; β_Vuc_ = 0.29 ± 0.07, p = 2·10^−4^), with no significant difference between the two (p = 0.36), therefore showing no evidence for a representation of the difference. We completed this simple analysis by a comparison using Bayesian Model Selection at the group level, between two variants of the main GLM where Val was replaced by either the sum (V_c_ + V_uc_) or the difference (V_c_ − V_uc_), competing to explain choice-related activity in a vmPFC ROI defined from the literature (to avoid non-independence issues). Although not formally conclusive, the comparison showed that exceedance probability was in favor of the sum model (Fig. S4D), thus validating our Val proxy as most relevant to capture vmPFC activity, even during choices. Another advantage of this Val proxy is being orthogonal to confidence, whereas the difference between option values is not. The consequence is that the neural correlates of Conf were unaffected by introducing the Val regressor, or by serial orthogonalization (Fig. S2).

Importantly, no consistent association with reward value or effort cost was observed in putative opponent brain regions such as the dmPFC, which was instead systematically reflecting DT. Thus, it appeared that dmPFC activity reflected the metacognitive effort cost invested in the ongoing task (deliberation about the response) rather than the effort cost attached to the option on valuation. Importantly, the association with DT was observed despite the fact that DT was orthogonalized to both value and confidence, suggesting that the dmPFC represents the effort invested above and beyond that induced by the difficulty of value-based judgment or decision. The parametric modulation by DT was also obtained when dmPFC activation was fitted with a boxcar function extending from stimulus response (Fig. S2), suggesting a modulation in amplitude beyond prolonged activity.

However, DT is a very indirect proxy for the effort invested in solving the task, and could be affected by many other factors (such as distraction or mind-wandering). We therefore investigated the relationship between brain activity and another proxy that has been repeatedly related to effort: pupil size. Neural activity was extracted in each ROI by fitting a GLM containing one event (stimulus onset) per trial. Then pupil size at each time point was regressed across trials against a GLM that contained factors of no interest (luminance, jitter duration, text length), variables of interest (Val, Conf, DT) and neural activity (vmPFC, mPFC, dmPFC).

A positive association between pupil size and dmPFC activity was observed in both rating and choice tasks (Fig. 4), about one second before the response. This association was not an artifact of the trial being prolonged (and therefore of the response to luminance being cut at different durations), since it was observed both when locking time courses on stimulus onset and on motor response (button press). Finally, it was specific to the dmPFC ROI, and observed even if dmPFC was made independent (through serial orthogonalization) to all other variables (notably Val, Conf and DT). Thus, the association between dmPFC and pupil size was observed above and beyond DT and factors that could affect DT. In contrast, there was no consistent association between vmPFC and pupil size before the response, suggesting that the correlates of DT observed in vmPFC were not related to effort but to some other factors affecting DT, such as mind-wandering.

**Figure 4:**
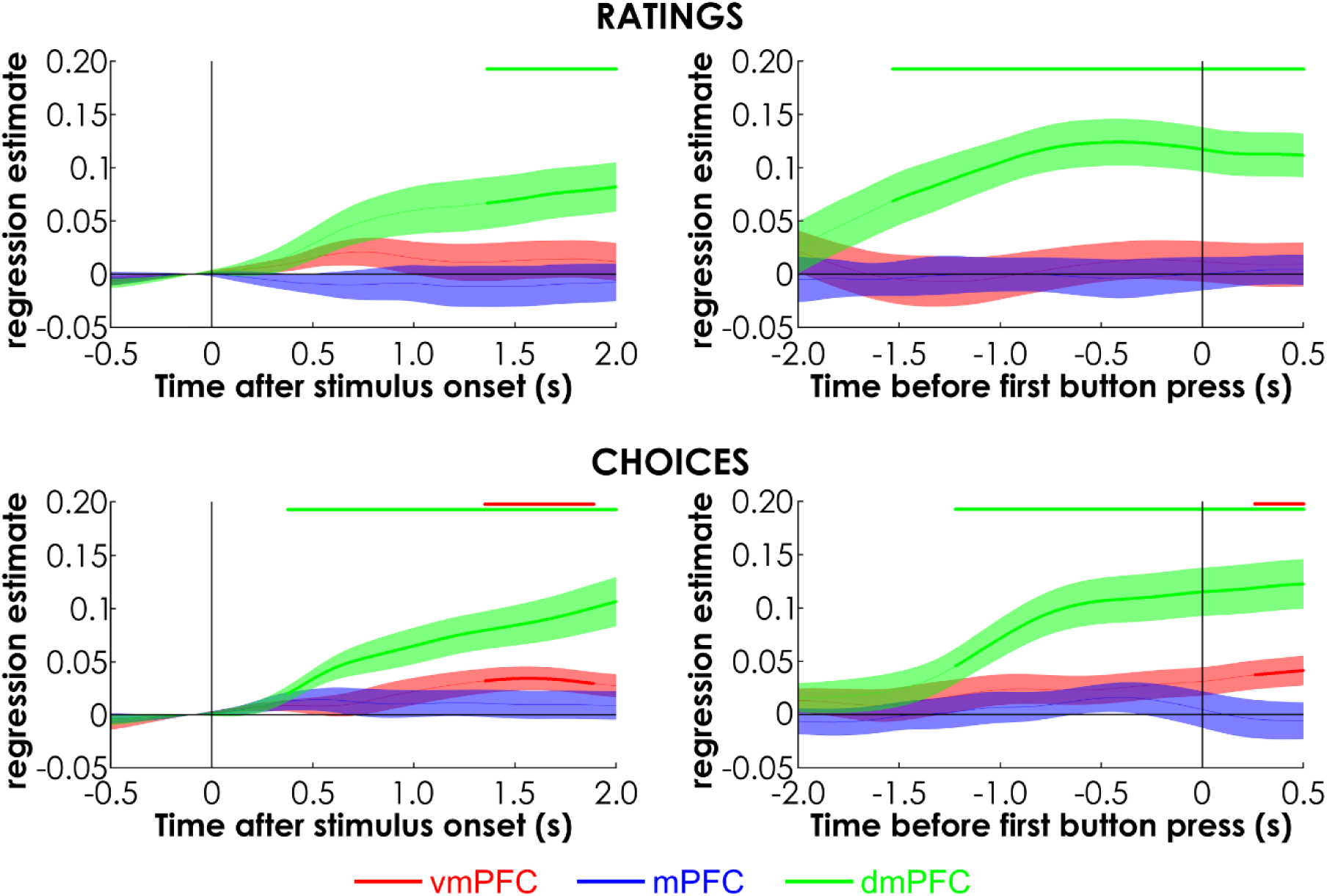
Pupillometric results. Plots show the time course of regression estimates, obtained with a GLM built to explain pupil size. The GLM included factors of no interest (jitter duration, stimulus luminance, text length), variables of interest (Val, Conf, DT) and activities in main ROI (vmPFC, mPFC, dmPFC, corresponding to red, blue and green traces, respectively). Each row corresponds to a different task (likeability rating, choice tasks). Left and right columns show time courses aligned onto stimulus onset and button press, respectively. Lines represent means across participants and shaded areas inter-participant standard errors. Horizontal bars indicate significant time clusters after correction for multiple comparisons using random-field theory.

## Discussion

Exploring the neural correlates of variables that are common to different preference expressing tasks, we observed a functional partition within the medial PFC: stimulus value, response confidence and deliberation time were best reflected in vmPFC, mPFC and dmPFC activity, respectively. These associations between regions and variables were consistent across rating and choice tasks, whether they involved likeability judgment, attribute integration or option comparison.

Our results confirm the role attributed to the vmPFC as a generic valuation system (Bartra et al., 2013; Levy and Glimcher, 2012). The subjective value of reward items was reflected in vmPFC activity irrespective of the category (food versus goods), as was reported in many studies (Abitbol et al., 2015; Chib et al., 2009; Lebreton et al., 2009; Lopez-Persem et al., 2020). Also, vmPFC value signals were observed whether or not reward items were presented with images, suggesting that they can be extracted from both direct perceptual input or from text-based imagination which was shown to recruit episodic memory systems (Lebreton et al., 2013). Critically, our results show that the vmPFC also reflects the effort cost (whether mental or physical) attached to potential courses of actions. Therefore, they challenge previous suggestions that the vmPFC is involved in stimulus valuation, independently of action costs (Pessiglione et al., 2018; Rangel and Hare, 2010). They rather suggest that the vmPFC might compute a net value, its activity increasing with reward benefit and decreasing with effort cost, so as to prescribe whether or not an action is worth engaging. This idea is in line with early demonstrations that the vmPFC integrates costs such as potential loss or delay in reward delivery (Hare et al., 2009; Kable and Glimcher, 2007; Talmi et al., 2009; Tom et al., 2007) and with recent mounting evidence that it also integrates effort-related costs (Aridan et al., 2019; Hogan et al., 2019; Lopez-Gamundi et al., 2021; Westbrook et al., 2019).

The other medial prefrontal clusters (mPFC and dmPFC) were not affected by reward values or effort costs attached to choice options, but by metacognitive variables related to expressing a preference, i.e. confidence and deliberation. Note that during a binary choice, confidence and deliberation are both related to the value difference between chosen and unchosen options. Here, response confidence (defined as the squared distance from the mean response) was orthogonal to stimulus value (defined as the addition of reward and/or effort values). The confidence proxy was previously shown to be highly correlated with confidence ratings and to elicit similar neural correlates (De Martino et al., 2017; Lopez-Persem et al., 2020). The value proxy is related to the representation of overall value (or ‘set liking’) assigned to choice options, which was previously observed in vmPFC activity (Blair et al., 2006; Hare et al., 2011; Jocham et al., 2014; Palminteri et al., 2009; Shenhav and Karmarkar, 2019). The two notions are close to the sum and difference signals that may emerge from an attractor network model in which two neuronal populations compete for their favorite option through mutual inhibition (Hunt et al., 2012). Our results suggest a dissociation of these signals, consistent with a previously described ventro-dorsal gradient from value to confidence (De Martino et al., 2017). The same dissociation applied to the rating task, where there is no comparison between unrelated options. We acknowledge that in a sense, likeability ratings can be conceived as a choice, since one position on the rating scale must be selected. However, this would be choosing between a large number (virtually infinite) of possible responses ordered along a single dimension (likeability). It is highly unlikely that the brain would solve the rating task through a competition mechanism in which each neuronal population would vote for one position on the scale. Thus, observing the same pattern of medial PFC activity across rating and choice tasks suggests that the functional role of this region cannot be reduced to models narrowly applied to the classical case of binary choice. However, rating and choice tasks both involve valuating the stimuli and selecting the response in which confidence is maximal, hence a common representation of stimulus value and response confidence in vmPFC and mPFC, respectively. Confidence was the only variable significantly associated to mPFC activity, but was also positively reflected in vmPFC activity, as previously reported (Chua et al., 2006; De Martino et al., 2013; Gherman and Philiastides, 2018). Indeed, the addition of value and confidence signals in the vmPFC is a pattern that has been already observed in both fMRI and iEEG activity (Lebreton et al., 2015; Lopez-Persem et al., 2020). On the contrary, dmPFC activity tended to decrease with confidence, but this trend did not survive significance threshold.

The variable that was robustly associated with dmPFC activity was deliberation time. This variable was not orthogonal to the others, since it decreased with both stimulus value and response confidence. In some of our analyses, deliberation time was post-hoc orthogonalized with respect to the other variables, meaning that the association with dmPFC activity was observed above and beyond the variance explained by stimulus value and response confidence. This association alone would not yield a clear-cut interpretation, since many factors may affect response time. However, the systematic link observed between trial-wise dmPFC activation and the increase in pupil size just before the response hints that this association might reflect the cognitive effort invested in the task. Indeed, pupil size has been associated to the intensity of not only physical effort, such as handgrip squeeze (Zénon et al., 2014) but also mental effort, such as focusing attention to resolve conflict or overcome task difficulty (Alnaes et al., 2014; Kahneman and Beatty, 1966; van der Wel and van Steenbergen, 2018). By contrast, we did not observe this systematic link with pupil size during deliberation with vmPFC activity. The link between vmPFC and deliberation time might therefore reflect other sources of variance, such as mind-wandering (being slower because of some off-task periods), in accordance with a previous report that elevated baseline vmPFC activity predicts prolonged response time (Hinds et al., 2013). Regarding dmPFC, our ROI overlaps with clusters that have been labeled dorsal anterior cingulate cortex, or sometimes pre-supplementary motor area, in previous studies (Kamiński et al., 2017; Kolling et al., 2016; Shenhav et al., 2013). The association with deliberation time is compatible with a role attributed to this region in the exertion of both physical effort (Klein-Flugge et al., 2016; Kurniawan et al., 2013; Skvortsova et al., 2014) and cognitive control (Botvinick et al., 2001; Kerns et al., 2004; Sohn et al., 2007).

To recapitulate, we have teased apart the neural correlates of likeability, confidence and deliberation in the medial prefrontal cortex, which have been confused in previous fMRI studies, as shown by meta-analytic maps. The key distinction operated here is perhaps between effort as an attribute of choice option and effort as a resource allocated to solving the task, or in other words, between valuation applied to effort (implicating the vmPFC) and effort invested in valuation (implicating the dmPFC). This dissociation is consistent with the idea that the vmPFC anticipates the aversive value of a potential effort, while the dmPFC represents the intensity of effort when it must be exerted. It could be related to efforts being hypothetical in our design, but previous studies have observed similar effort representation in the vmPFC (not the dmPFC) when efforts were not hypothetical but only implemented later, at the end of the experiment (Aridan et al., 2019; Hogan et al., 2019; Westbrook et al., 2019). At a metacognitive level, our results could be interpreted in the frame of a resource allocation model, where the effort or time invested in the deliberation is meant to increase confidence in the response, whether a rating or a choice (Lee and Daunizeau, 2021). Yet our results cannot tell whether the dmPFC signals the need for deliberation effort, monitors the time invested in deliberation, or generates an aversive feeling related to the prolongation of deliberation.

Even if showing robust associations between brain regions and cognitive variables, our approach (looking for robust associations across tasks) also bears limitations. Notably, our design would not allow comparing between conditions, as is traditionally done in neuroimaging studies. One may want for instance to compare between tasks and test whether brain regions are more involved in one or the other, but this would be confounded by several factors, such as the order (choice tasks being performed after rating tasks). A significant contrast would not be interpretable anyway, because there is more than one minimal difference between tasks. Thus, the aim to generalize the role of brain regions across tasks carries the inherent drawback of a limited specificity, but also the promises of a more robust understanding of anatomo-functional relationships. We hope this study will pave the way to further investigations following a similar approach, assessing a same concept across several tasks in a single study, instead of splitting tasks over separate reports, with likely inconsistent conclusions.

## Methods

### Subjects

In total, 40 right-handed volunteers participated in this fMRI study, which was approved by the Pitié-Salpêtrière Hospital local ethics committee. Participants were recruited through the RISC (Relais d’Information en Sciences de la Cognition) online platform (https://www.risc.cnrs.fr/) and signed inform consent prior to participation in the study. All participants were screened for the use of psychotropic medications and drugs, history of psychiatric and neurologic disorders, and traumatic brain injury. One participant was excluded from all analyses because of a clear misunderstanding about task instructions, leaving n=39 participants for behavioral data analysis (22 females / 17 males, aged 25.4±4.1 years). Another participant was excluded from the fMRI analysis due to excessive movement inside the scanner (>3mm within-session per direction). Eleven additional participants were excluded from pupil size analysis, due to poor signal detection in at least one of the sessions (leaving a total of n=27 participants for pupil analysis).

All participants gave informed consent and were paid a fixed amount for their participation. The 15 first subjects were paid 60€ and the 25 other subjects were paid 75€. The difference in payoff corresponds to a difference in scanning protocols, although all participants performed the same tasks. The pilot protocol (n=15) aimed at comparing fMRI data acquisition sequences: regular EPI, EPI with multiband acceleration, EPI with multiband acceleration + multi-echo acquisition. The main protocol (n=25) aimed at addressing the neurocognitive question of interest with the best acquisition sequence. For this main protocol, we kept the regular EPI sequence for all sessions, as we saw no clear advantage for multiband acceleration or multi-echo acquisition in basic contrast images. Therefore, the analyses only include fMRI data using regular EPI acquisition (three sessions for the pilot protocol, all nine sessions for the main protocol).

### Behavioral tasks

All tasks were programmed using the Psychtoolbox (Brainard, 1997) Psychtoolbox-3 running in Matlab (The MathWorks, Inc., Version 2012). Participants were given a 4-button box (fORP 932, Current Designs Inc, Philadelphia, USA) placed under their right hand to provide their responses. Stimuli were projected on a computer screen, their luminance being estimated using standard function of red-green-blue composition (0.299·red + 0.587·green + 0.114·blue, see http://www.w3.org/TR/AERT#color-contrast). Stimuli comprised 144 reward items (72 food and 72 goods) and 72 effort items (36 mental and 36 physical). Half the reward items were presented with text only (R_t_ items), and the other half was presented with both text and image (R_ti_ items).

All effort items were only described with text (E_t_). For each task, fMRI sessions were preceded by a short training (not included in the analysis), for participants to familiarize with the sort of items they would have to valuate and with the button pad they would use to express their preferences.

Participants all started with a (dis-)likeability rating task (Fig. 1A), performed during the first three fMRI sessions, each divided into three 24-trial blocks corresponding to the three stimulus type (R_ti_, R_t_, E_t_). The order of blocks within a session was counterbalanced across participants. The items were presented one by one, and participants rated them by moving a cursor along a visual analog scale. They used their index and middle fingers to press buttons corresponding to left and right movements, and validated the final position of the cursor by pressing a third button, which triggered the new trial. The initial position of the cursor, at the beginning of each trial, was randomly placed between 25 and 75% of the 0-100 rating scale. There was no mark on the scale, giving the impression of a continuous rating, although it was in practice discretized into 100 steps. The left and right extremes of the scale were labeled “I would not care” and “I would like it enormously” for reward items, “I would not mind” and “I would dislike it enormously” for effort items. Note that both reward and effort scales included indifference at one extremity, such that the two scales could form a continuum of increasing likeability from very aversive effort to very appetitive reward. In any case, the situations to be rated were hypothetical: the question was about how much they would like the reward (should it be given to them at the end of the experiment) and how much they would dislike the effort (should it be imposed to them at the end of the experiment). Should the timeout (10 s in rating tasks and 5s in choice tasks) be reached, the message ‘too slow’ would have been displayed on screen and the trial repeated later, but this remained exceptional.

After the three rating sessions, participants performed a series of binary choices. The A/B choice task (Fig. 1B) involved expressing a preference between two options of a same dimension, presented on the left and right of the screen. The two options were items presented in the rating task, drawn from the same category, regarding both the presentation mode (R_ti_ vs R_ti_, R_t_ vs R_t_, E_t_ vs E_t_) and type of stimulus (food vs. food, goods vs. goods, mental vs mental, physical vs physical). Each item was presented twice, following two intermixed pairing schedules: one varied the mean rating (i.e., stimulus value) while controlling for distance (i.e., decision value or choice difficulty), whereas the other varied the distance in rating while controlling the mean. Participants selected the reward they would most like to obtain, or the effort they would least dislike to exert, by pressing the left or right button with their middle or index finger. The chosen option was then highlighted with a red frame, so participants could check that their choice was correctly recorded. The fMRI sessions devoted to the A/B choice task included three 24-trial blocks presenting the three types of options (R_ti_, R_t_, E_t_), the order of blocks being counterbalanced across participants.

Then participants performed the Yes/No choice task (Fig. 1C), which involved deciding whether to accept exerting a given effort in order to get a given reward. Thus, every trial proposed one option combining two dimensions (one R_t_ and one E_t_ item). Each item was presented twice, following two intermixed pairing schedules: one associating more pleasant reward with more painful effort (thus controlling for decision value or choice difficulty), the other associating more pleasant reward with less painful effort (thus varying choice difficulty). The mean net value was also balanced across fMRI sessions. Participants selected their response by pressing the button corresponding to ‘yes’ or ‘no’ with their index or middle finger. The left/right position of yes/no responses was counterbalanced across trials. To give participants a feedback on their choice, the selected option was highlighted with a red frame. The three fMRI sessions devoted to the Yes/No choice task contained 48 trials each.

Note that, as were ratings, all choices were hypothetical. This was implemented to enable the use of natural reward and effort items that can be encountered in everyday life but are difficult to implement in the lab (such as walking a 1-km distance). Another reason was to allow for a distinction between the estimation of effort cost and motor preparation processes that are triggered when efforts are implemented (Hogan et al., 2019).

### Behavioral data analysis

All data were analyzed using Matlab 2017a (The MathWorks, Inc., USA). Choices were fitted with logistic regression models of decision value, with intercept and slope parameters.

For A/B choices, the model was:

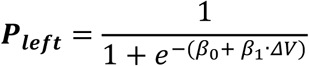

Where P_left_ is the probability of choosing the left option, ΔV is the decision value, i.e. the difference in likeability rating between left and right options (V_left_ - V_right_), while β_0_ and β_1_ are the intercept and slope parameters capturing potential bias and choice consistency (inverse temperature).

For Yes/No choices, the model was:

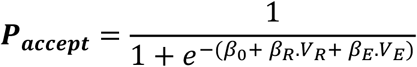

Where P_accept_ is the probability of accepting the offer (make the effort to get the reward), V_R_ and V_E_ are the likeability ratings provided for the reward and effort items. Thus, the decision value (or net value) here is a weighted sum of reward and effort likeability (one being positive and the other negative), the parameter weights β_R_ and β_E_ serving as both scaling factors and inverse temperature.

The stimulus value (Val) and response confidence (Conf) regressors used in the analysis of deliberation time (DT) and fMRI data were respectively defined as the addition of likeability ratings assigned to the items on screen and the squared distance from the mean response. They were adapted to each task, as follows:

|  | Rating task | A/B choice task | Yes/No choice task |
| --- | --- | --- | --- |
| Val | V | $V_{\text{left}} + V_{\text{right}}$ | $\beta_R \cdot V_R + \beta_E \cdot V_E$ |
| Conf | $[V - \text{mean}(V)]^2$ | $[P_{\text{left}} - \text{mean}(P_{\text{left}})]^2$ | $[P_{\text{accept}} - \text{mean}(P_{\text{accept}})]^2$ |

In each case, V is either the reward or effort likeability provided by z-scored individual rating of the item presented in a given trial, and P is the probability generated for each trial using the logistic model fitted to choices. Note that, by construction (before z-scoring), V is positive for reward items (which are liked) and negative for effort items (which are disliked). The mean response is simply the mean rating over trials, the mean frequency of left choice and the mean frequency of accept choice, depending on the task.

Deliberation time (DT) was defined across tasks as the time between stimulus onset and first button press. Trial-wise variations in DT were fitted with linear regression models, including a session-specific intercept, factors of no interest - fixation cross, display duration (Jitter), stimulus luminance (Lum), text length in number of words (Length) - and factors of interest - stimulus value (Val), response confidence (Conf). Thus, the model was:

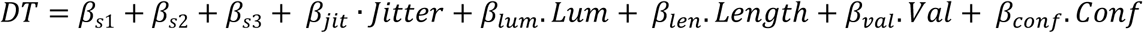

### fMRI data acquisition

Functional and structural brain imaging data was collected using a Siemens Magnetom Prisma 3-T scanner equipped with a Siemens 64 channel Head/Neck coil. Structural T1-weighted images were coregistered to the mean echo planar image (EPI), segmented and normalized to the standard T1 template and then averaged across subjects for anatomical localization of group-level functional activation. Functional T2*-weighted EPIs were acquired with BOLD contrast using the following parameters: repetition time TR = 2.01 seconds, echo time TE = 25 ms, flip angle = 78°, number of slices = 37, slice thickness = 2.5 mm, field of view = 200 mm. A tilted-plane acquisition sequence was used to optimize sensitivity to BOLD signal in the orbitofrontal cortex (44). Note that the number of volumes per session was not predefined, because all responses were self-paced. Volume acquisition was just stopped when the task was completed.

Most subjects (n=25) performed nine fMRI sessions (three per task) using this standard EPI sequence. The pilot subgroup (n=15) also performed nine fMRI sessions, but the fMRI data acquisition sequences were alternated between standard EPI, EPI with multi-band acceleration factor (TR = 1.20 s; TE = 25 ms; flip angle = 66°; number of slices = 44; slice thickness = 2.5 mm; acceleration factor = 2) and EPI with multi-band acceleration factor + multi-echo (TR = 1.28 s; TE = 11.00 ms and 29.89 ms; flip angle = 69°; number of slices = 44; slice thickness = 2.5 mm; acceleration factor = 2). The order of fMRI sequences was counterbalanced across participants. Preliminary analyses of basic contrast images were done using the pilot dataset to select the best acquisition sequence. As there was no clear benefit with the multi-band and multi-echo add-ons, we retained the standard EPI for the main experiment. Results obtained with the other fMRI sequences are shown in extended figures (Fig. S3).

### fMRI data analysis

Functional MRI data were preprocessed and analyzed with the SPM12 toolbox (Wellcome Trust Center for NeuroImaging, London, UK) running in Matlab 2017a. Preprocessing consisted of spatial realignment, normalization using the same transformation as anatomical images, and spatial smoothing using a Gaussian kernel with a full width at a half-maximum of 8 mm.

Preprocessed data were analyzed with a standard general linear model (GLM) approach at the first (individual) level and then tested for significance at the second (group) level. All GLM included the six movement regressors generated during realignment of successive scans. In our main GLM, stimulus onset was modeled by a stick function, modulated by the following regressors: 1) fixation cross duration, 2) luminance, 3) text length, 4) Val, 5) Conf, 6) DT. The first three were factors of no interest that were found to significantly impact DT in the linear regression analysis. The regressors of interest (Val, Conf and DT) were defined as explained in the behavioral data analysis section. The different blocks of the rating and A/B choice tasks (presenting reward as text + image, reward as text and effort as text) were modeled in separate regressors. All regressors of interest were z-scored and convolved with the canonical hemodynamic response function and its first temporal derivative. All parametric modulators were serially orthogonalized. At the second level, correlates of Val, Conf and DT were obtained with contrasts tested across tasks of corresponding regression estimates against zero. Note that likeability ratings obtained for effort items were negative in all regressors (meaning that they can only decrease stimulus value).

Several alternative GLM were built to test variants of the main GLM. GLM2 was identical to GLM1 except that orthogonalization was removed such that all native regressors could compete to explain variance in fMRI time series. GLM3 was identical to GLM1, except that instead of a stick function, stimulus onsets were modeled with a boxcar function modeling periods from stimulus onset to first button press. Three additional GLM were built to further explore the choice tasks. In GLM4, the Val regressor (sum of option values) was replaced by the difference between option values (V_c_ − V_uc_) for the two choice tasks. This GLM served to perform a group-level Bayesian model comparison to test which value regressor (sum or difference) best explains the fMRI time series during choice tasks. In GLM5, Conf and DT were removed and Val was replaced by two separate regressors for the chosen and unchosen option values (V_c_ and V_uc_). This GLM was used to test whether regressor estimates for chosen and unchosen values had the same sign (as in a sum) or opposite signs (as in a difference). In GLM6, reward and effort values were split in two separate regressors for all tasks (including the Yes/No choice task). The purpose of this GLM was to distinguish between neural correlates of reward value and effort cost in brain valuation regions. Finally, a last GLM was built with one event per trial, modeled with a stick function, at the time of stimulus onset, with the aim to extract trial-by-trial activity levels in regions of interest, which then served as regressors to explain pupil size data (see next section).

Regions of interest (ROI) were defined as clusters in group-level statistical maps that survived significance threshold of p < 0.05 after family-wise error correction for multiple comparisons at the voxel level. To avoid double dipping (Kriegeskorte et al., 2009) in statistical tests, regression estimates were extracted from ROI re-defined for each participant through a leave-one-out procedure. Regarding Bayesian Model Selection, to avoid biasing the comparison in favor of one or the other GLM, an independent ROI was defined as the conjunction between the positive minus negative value contrast in a published meta-analysis (Bartra et al., 2013) and the bilateral medial orbitofrontal cortex region from the AAL atlas (Tzourio-Mazoyer et al., 2002). Additionally, we defined twelve 8-mm radius spherical ROI in the medial wall to illustrate the distribution of regression estimates for Val, Conf and DT. Parameter estimates were extracted from each voxel within these ROI and then averaged across voxels.

### Meta-analysis of fMRI studies

The meta-analytic maps were extracted from the online platform Neurosynth (https://www.neurosynth.org/), using the keywords “value” (470 studies), “confidence” (79 studies) and “effort” (204 studies) for “uniformity test”, which displays brain regions that are consistently activated across papers mentioning the target keyword. Each map was binarized to visualize clusters surviving a significance threshold of p < 0.01 after false discovery rate (FDR) correction for multiple comparisons.

### Pupil size

Pupil diameter was recorded at a sampling rate of 1000Hz, using an EyeLink 1000 plus (SR Research) eye-tracker. The eye-tracker was calibrated before the start of fMRI sessions, once the subject was positioned inside the scanner. A cubicle interpolation was performed to compensate for any period of time when the pupil signal was lost due to blinking. The pupil size time series were subsequently band-pass filtered (1/128 to 1 Hz) and z-scored per session.

Within-trial variations in pupil size was baseline-corrected (by removing the mean signal over the 200 ms preceding stimulus onset) and time-locked either to stimulus onset or button press. Then trial-wise variations in pupil size were fitted with a linear regression model that included factors of no interest (an intercept per block, jitter duration, stimulus luminance and text length), variables of interest (Val, Conf and DT defined as in the behavioral data analysis section) and neural activity (extracted from vmPFC, mPFC and dmPFC ROI clusters). Within-trial individual time series of regression estimates were then smoothed using a 100ms kernel. Group-level significant time clusters were identified after correction for multiple comparisons estimated according to random field theory, using the RFT_GLM_contrast.m function of the VBA toolbox (available at http://mbb-team.github.io/VBA-toolbox/).

## Acknowledgments

We would like to thank the CENIR (research neuroimaging center) staff for their help in fMRI data acquisition (particularly Stéphane Lehéricy, Romain Valabrègue and Mathieu Santin for the optimization of scanning sequences), Chen Hu for assistance in data collection, Jules Brochard for assistance in data analysis, Fabien Vinckier and Jean Daunizeau for insightful comments. The study was funded by a research grant from the “Fondation pour la Recherche Médicale” and by the “Investissements d’avenir” program (ANR-10-IBHU-0003).

