## Supplementary Material for "Value, confidence, deliberation: a functional partition of the medial prefrontal cortex demonstrated across rating and choice tasks"

#### Supplementary Figures

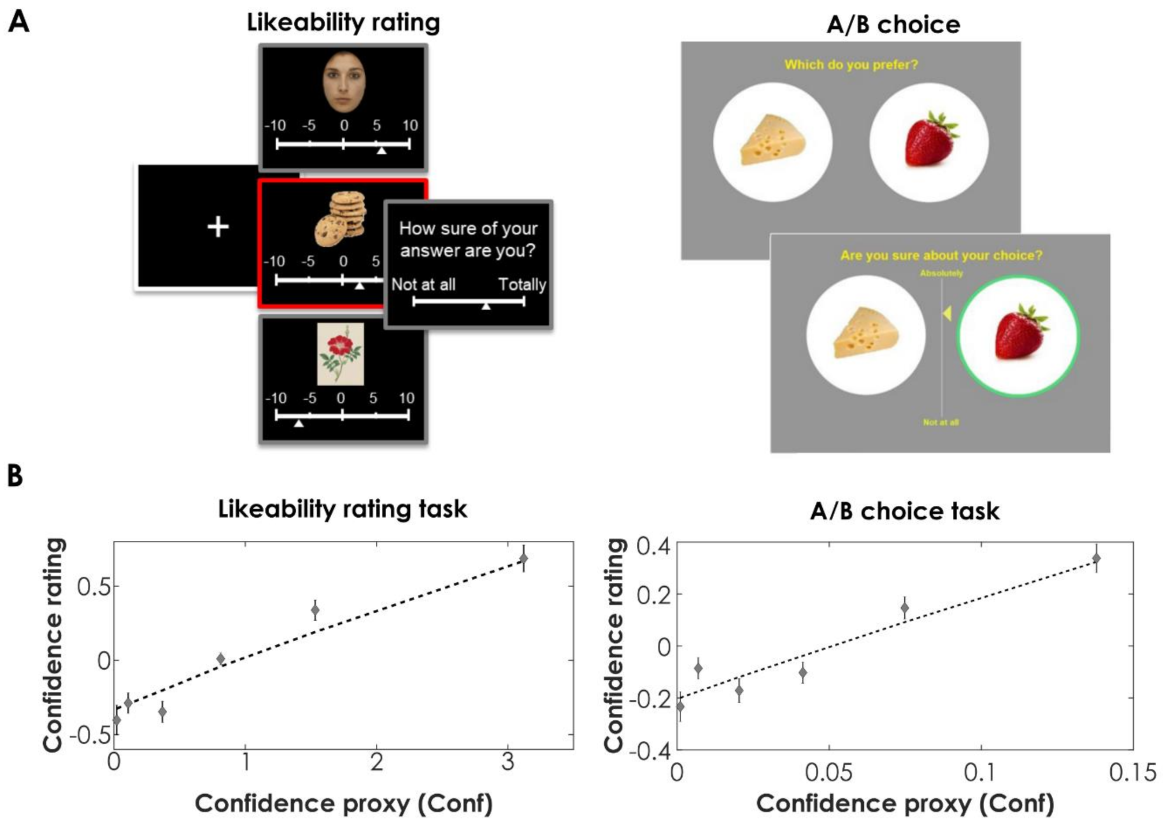


**Figure S1: Testing the validity of our proxy for confidence (Conf) in other datasets.**

**A]** In the likeability rating task (Lopez-Persem et al., 2020), participants first rated the likeability of food, face and painting items and then provided a confidence rating of their own likeability judgment. In the 1D-20 binary choice task (Lee and Daunizeau, 2020), participants selected their preferred item between options shown in pairs, and then provided a confidence rating about having made the best choice.

B] The graphs show confidence rating as a function of binned Conf (square of centered likeability rating or centered choice likelihood). Dots represent mean over participants, error bars are inter-participant standard errors, dotted lines show linear regression fits.

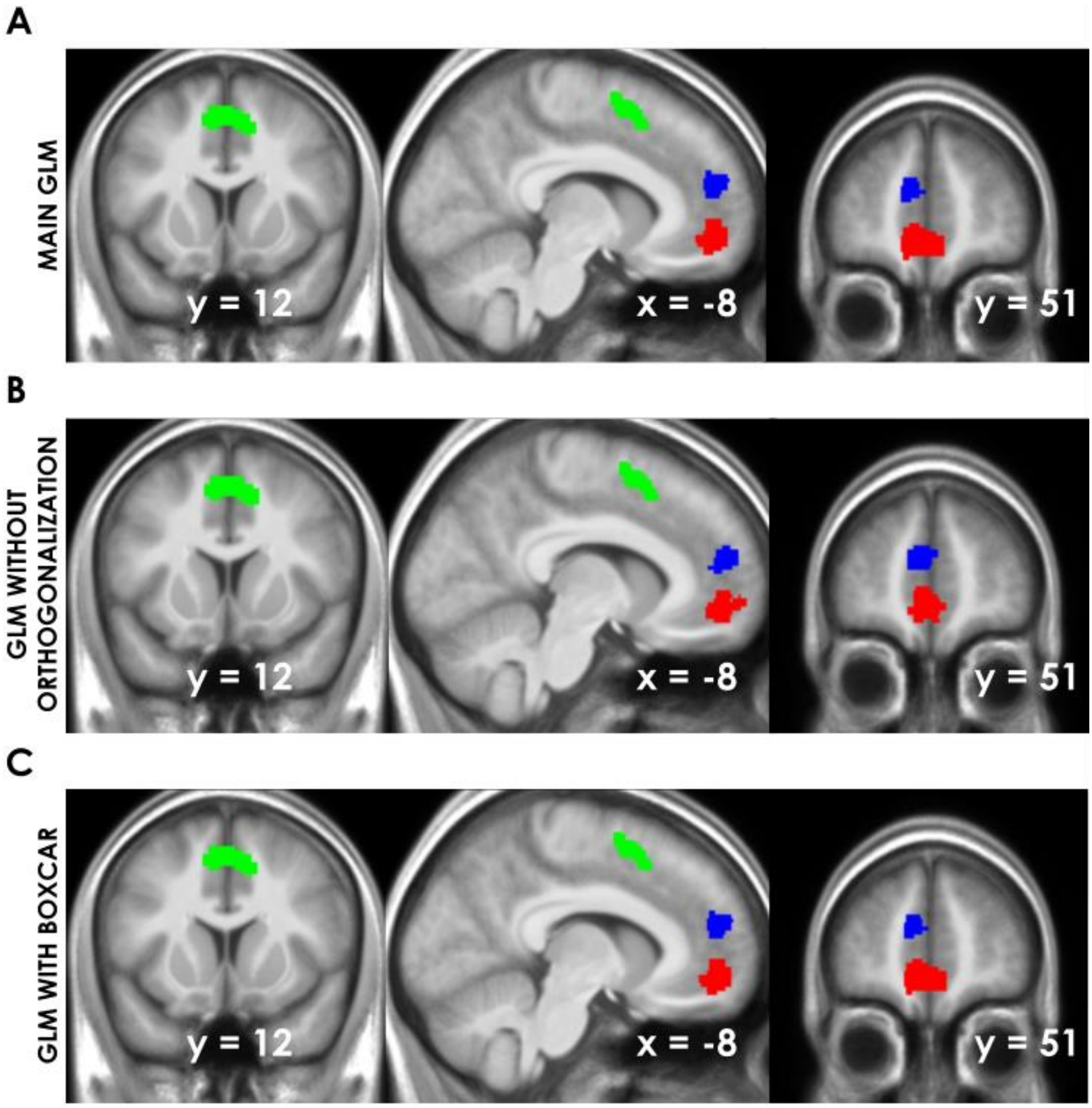


**Figure S2: Neural representations of value, confidence and deliberation constructs in the medial prefrontal cortex obtained with alternative GLM.**

A] Statistical map (same as in Fig. 3B) obtained with the main GLM is shown for comparison. B] Statistical map obtained with the same GLM when serial orthogonalization was removed. C] Statistical map obtained with the same GLM when events were modeled with a boxcar function encompassing the period from trial onset to first button press. For all maps, sagittal slices were taken at the same coordinates as the Neurosynth output (shown in Fig. 3A), and superimposed on the average anatomical scan normalized to canonical (MNI) template. Maps were thresholded at $p < 0.05$ after voxel-wise family-wise error correction for multiple comparisons. For all maps, only the main clusters of interest located in the medial prefrontal cortex are shown. For clusters outside the medial prefrontal cortex, please refer to Tables S4-6.

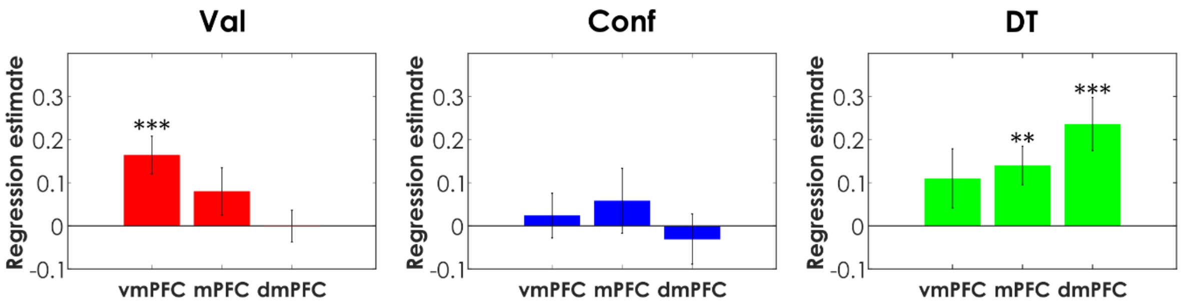


**Figure S3: Partial replication in independent smaller dataset (n=15) using different fMRI sequences.**

Data were collected in a subset of participants and task sessions scanned with different fMRI data acquisition sequences. Regression estimates obtained for value, confidence and deliberation were extracted from 8mm-radius spheres centered on activation peaks in group-level statistical maps of activity recorded with standard EPI sequence (clusters shown in Fig. 3B). The bars correspond to the average across the two other sequences used, namely one EPI sequence with a multiband acceleration factor and one multi-echo sequence with a multiband acceleration factor. Bars show mean across participants, error bars show inter-participant standard errors. Stars indicate significance of t-test against zero (*** $p < 0.005$, ** $p < 0.01$, * $p < 0.05$). The trends were similar to those observed with the standard EPI sequence, but were not always significant due to low statistical power.

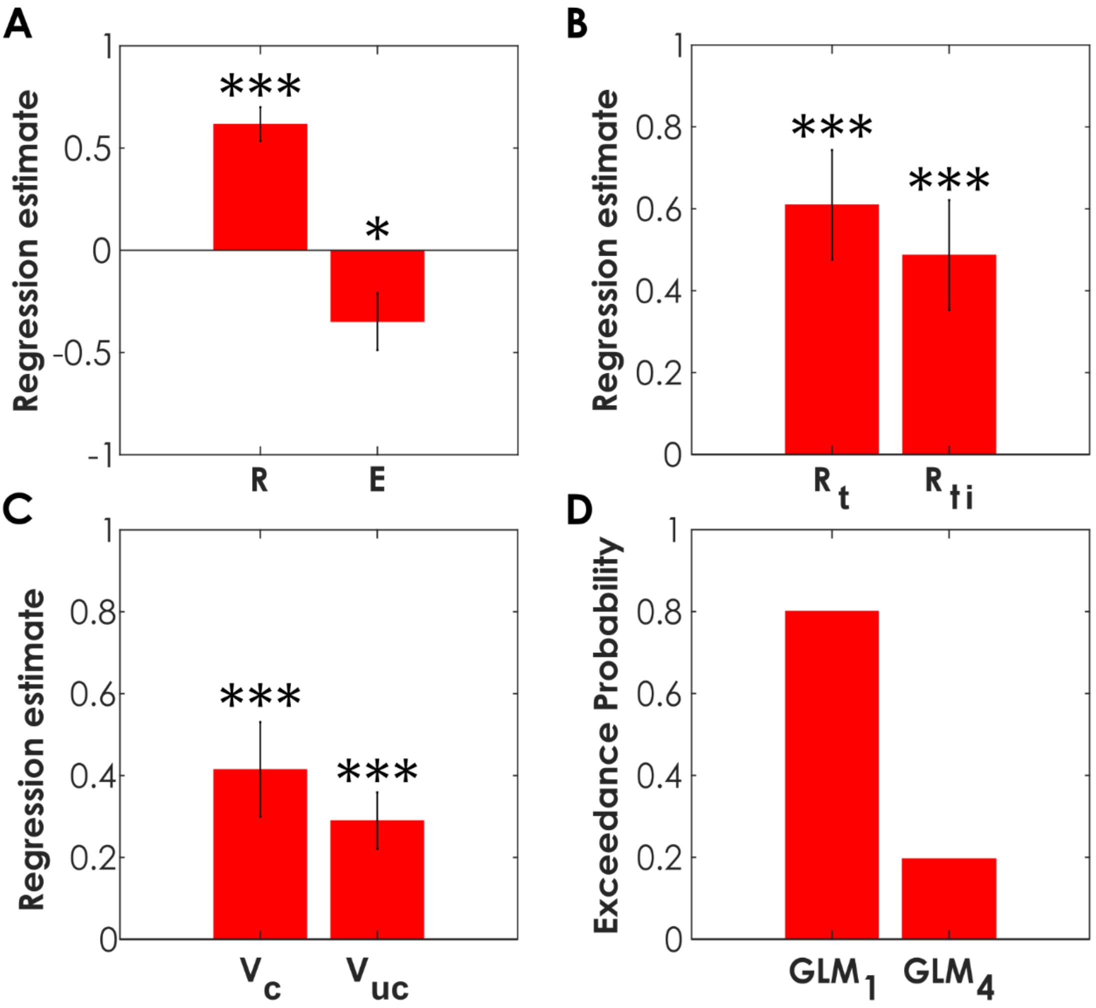


**Figure S4: Further specification of value-related activity in the vmPFC**

A and B] Regression estimates were extracted across rating and choice tasks, separately for rewards presented as text (Rt) or text + image (Rti) and separately for reward (R) and effort (E) values (results in A are from GLM6 where reward and effort regressors are split even for the Yes/No choice task, while results in B are from GLM1). The vmPFC ROI was based on group-level cluster activated with Val using GLM1, following a leave-one out procedure to avoid double dipping.

C] Regression estimates were extracted from the vmPFC (group-level cluster associated to Val), using GLM5 where Val, Conf and DT were replaced by the chosen and unchosen option values ($V_c$ and $V_{uc}$), across the two choice tasks. In more details, $V_c$ / $V_{uc}$ were $V_{left}$ / $V_{right}$ for a left choice in the A/B task, and $\beta_R \cdot V_R + \beta_E \cdot V_E$ / 0 for a yes choice in the Yes/No task (and vice-versa for opposite choices).

D] Results of a Bayesian Model Comparison between GLM1, where Val is the sum, and GLM4, where Val is the difference between option values ($V_c$ - $V_{uc}$), for explaining vmPFC activity across the two choice tasks. The vmPFC was defined by a conjunction between the correlates of positive minus

negative value from a published meta-analysis (Bartra et al., 2013) and the medial prefrontal cortex region from the AAL atlas (Tzourio-Mazoyer et al., 2002) to avoid biasing the comparison in favor of GLM1. Exceedance probability estimates were averaged across all voxels within the vmPFC ROI. Note that similar results were obtained when restricting the comparison to the A/B choice task.
Stars indicate significance of t-test against zero (*** $p < 0.005$, ** $p < 0.01$, * $p < 0.05$).

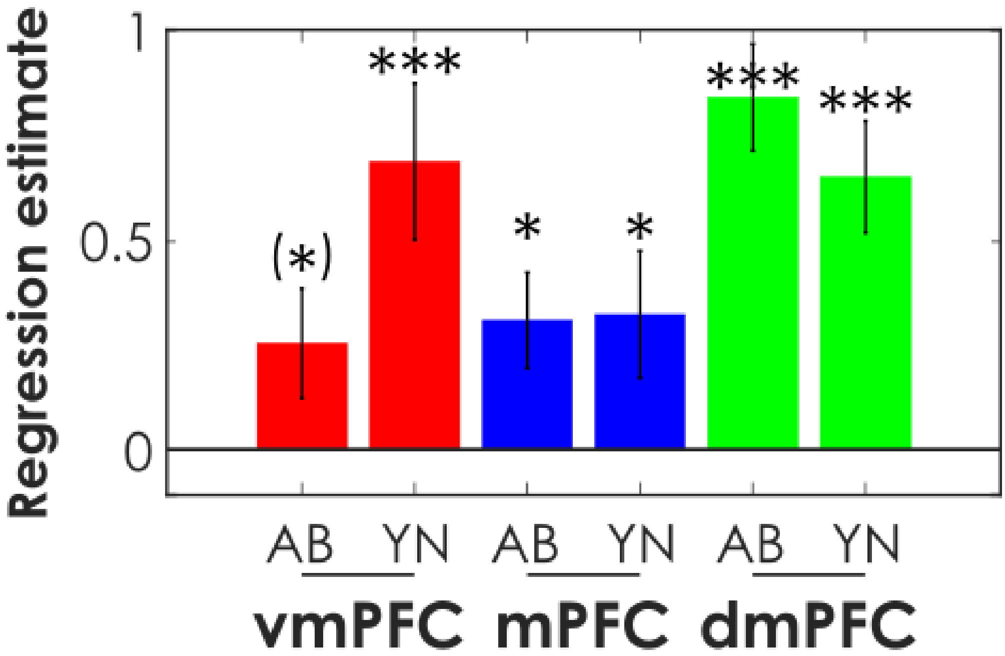


**Figure S5: Decomposition of regression estimates obtained for each variable of interest (Val, Conf and DT), plotted separately for each choice task (noted A/B and Y/N) in the different ROI (vmPFC, mPFC, dmPFC).**

The ROI have been selected with a leave-one out procedure as in the main figure (Fig. 3D).

Bars show means over participants, error bars show inter-participant standard errors. Stars indicate significance of t-test against zero (*** $p < 0.001$, ** $p < 0.01$, * $p< 0.05$, (*) $p < 0.10$). For the three region – variable associations, there was no significant difference between regression estimates obtained in the A/B and Yes/No choice tasks.

### Supplementary Tables

**Table S1: Brain activity signaling stimulus value (Val) across rating and choice tasks.**
Regions survived a significance threshold of $P < 0.05$ after FWE correction for multiple comparisons at the voxel level. Clusters smaller than 12 voxels, corresponding to the size of our smoothing kernel, were excluded from the table. Coordinates refer to the MNI space. The p-value reported is the p-value of the cluster after a FWE correction at the cluster level.

| Region | P cluster | Peak x | Peak y | Peak z | No. of Voxels |
|---|---|---|---|---|---|
| vmPFC | $3 \cdot 10^{-10}$ | -10 | 48 | -12 | 364 |
| Lingual Gyrus | $1 \cdot 10^{-4}$ | 16 | -70 | -6 | 64 |
| Orbitofrontal cortex | $2 \cdot 10^{-4}$ | -28 | 36 | -14 | 57 |
| Posterior cingulate cortex | 0.003 | -6 | -54 | 14 | 22 |
| Cingulate Gyrus | 0.005 | -8 | 38 | 6 | 16 |

**Table S2: Brain activity signaling response confidence (Conf) across rating and choice tasks.** Regions survived a significance threshold of $P < 0.05$ after FWE correction for multiple comparisons at the voxel level. Clusters smaller than 12 voxels, corresponding to the size of our smoothing kernel, were excluded from the table. Coordinates refer to the MNI space. The p-value reported is the p-value of the cluster after a FWE correction at the cluster level.

| Region | P FWE cluster | Peak x | Peak y | Peak z | No. of Voxels |
|---|---|---|---|---|---|
| mPFC | $5 \cdot 10^{-6}$ | -8 | 52 | 18 | 128 |
| Middle Temporal Gyrus | $2 \cdot 10^{-4}$ | -56 | -26 | -10 | 63 |
| Supramarginal Gyrus | $3 \cdot 10^{-4}$ | -62 | -40 | 32 | 56 |
| Middle Temporal Gyrus | 0.003 | -46 | -64 | 12 | 22 |
| Caudate Nucleus | 0.006 | -12 | 14 | -12 | 14 |
| Inferior Temporal Gyrus | 0.007 | -46 | 2 | -36 | 13 |

**Table S3: Brain activity signaling deliberation time (DT) across rating and choice tasks.** Regions survived a significance threshold of $P < 0.05$ after FWE correction for multiple comparisons at the voxel level. Clusters smaller than 12 voxels, corresponding to the size of our smoothing kernel, were excluded from the table. Coordinates refer to the MNI space. The p-value reported is the p-value of the cluster after a FWE correction at the cluster level.

| Region | P FWE cluster | x | y | z | No. of Voxels |
|---|---|---|---|---|---|
| dmPFC | $1 \cdot 10^{-9}$ | 10 | 12 | 48 | 365 |
| Inferior Frontal Gyrus | $7 \cdot 10^{-8}$ | -40 | 22 | 24 | 242 |
| Anterior Insula (left) | $2 \cdot 10^{-5}$ | -30 | 26 | 4 | 110 |
| Anterior Insula (right) | $4 \cdot 10^{-5}$ | 32 | 26 | 4 | 95 |
| Lingual Gyrus | 0.003 | -18 | -88 | -10 | 23 |

**Table S4: Brain activity signaling stimulus value (Val) across rating and choice tasks when regressors were not orthogonalized.**

Regions survived a significance threshold of $P < 0.05$ after FWE correction for multiple comparisons at the voxel level. Clusters smaller than 12 voxels, corresponding to the size of our smoothing kernel, were excluded from the table. Coordinates refer to the MNI space. The p-value reported is the p-value of the cluster after a FWE correction at the cluster level.

| Region | P cluster | Peak x | Peak y | Peak z | No. of Voxels |
|---|---|---|---|---|---|
| vmPFC | $1 \cdot 10^{-12}$ | -10 | 44 | -10 | 423 |
| Cingulate Gyrus | $2 \cdot 10^{-4}$ | -4 | 40 | 4 | 46 |
| Lingual Gyrus | 0.004 | -12 | -50 | 4 | 14 |

**Table S5: Brain activity signaling response confidence (Conf) across rating and choice tasks when regressors were not orthogonalized.**

Regions survived a significance threshold of P < 0.05 after FWE correction for multiple comparisons at the voxel level. Clusters smaller than 12 voxels, corresponding to the size of our smoothing kernel, were excluded from the table. Coordinates refer to the MNI space. The p-value reported is the p-value of the cluster after a FWE correction at the cluster level.

| Region | P FWE cluster | Peak x | Peak y | Peak z | No. of Voxels |
|---|---|---|---|---|---|
| mPFC | $1 \cdot 10^{-7}$ | -6 | 52 | 18 | 222 |
| Temporal mid pole | 0.001 | -60 | -28 | -10 | 47 |
| Temporal Superior Pole | 0.007 | -36 | 18 | -26 | 13 |
| Inferior Frontal Gyrus | 0.007 | -40 | 28 | -2 | 13 |

**Table S6: Brain activity signaling deliberation time (DT) across rating and choice tasks when regressors were not orthogonalized.**

Regions survived a significance threshold of $P < 0.05$ after FWE correction for multiple comparisons at the voxel level. Clusters smaller than 12 voxels, corresponding to the size of our smoothing kernel, were excluded from the table. Coordinates refer to the MNI space. The p-value reported is the p-value of the cluster after a FWE correction at the cluster level.

| Region | P FWE cluster | x | y | z | No. of Voxels |
|---|---|---|---|---|---|
| dmPFC | $8 \cdot 10^{-10}$ | 10 | 12 | 48 | 370 |
| Inferior Frontal Gyrus | $5 \cdot 10^{-8}$ | -40 | 22 | 24 | 249 |
| Anterior Insula (left) | $1 \cdot 10^{-5}$ | -30 | 26 | 4 | 110 |
| Anterior Insula (right) | $3 \cdot 10^{-5}$ | 32 | 26 | 4 | 96 |
| Lingual Gyrus | 0.003 | -18 | -88 | -10 | 23 |
